# Structural grammar of truncation across the human homodimer landscape

**DOI:** 10.64898/2026.05.06.723091

**Authors:** Taner Karagöl, Alper Karagöl

## Abstract

Alternative splicing and proteolytic truncation generate tens of thousands of protein isoforms in the human proteome. Their consequences for quaternary state, the level at which most protein function operates, have been examined largely one molecule at a time. Using the AlphaFold Protein Structure Database expansion to predicted human homodimers, we compared 5,168 canonical-versus-truncated pairs on four interface metrics distributed with that release. Truncation partitions high-confidence canonical homodimers into structural conservation (56.4% of pairs), complete interface ablation (26.1%) and partial destabilization (17.5%). The partition holds on predicted interface size and not only on interface confidence. Conservation events retain a median 0.938 of the canonical interface residues, destabilization events 0.622, and ablation events 0.000. Interface loss is graded by deletion magnitude, falling from 0.997 of the interface retained below a tenth of the chain removed to 0.192 above a half (Spearman rho = -0.552, p = 4.5 × 10^−61^). Deletion topology, derived by global alignment, is 53.3% N-terminal and 42.7% C-terminal. C-terminal truncation lowers interface conservation from 65.4% to 37.5%, an asymmetry confined to deletions above the cohort median of 89 residues. Of 208 pairs in which ipTM reports emergence from a sub-threshold canonical baseline, 42 reproduce on ipSAE, the metric on which the release defines its tiers. Canonical disorder content shows no detectable difference between emergence events and controls matched on canonical confidence. Predicted confidence tracks deposited structures, with two or more copies in a deposited assembly for 95.3% of high-confidence canonicals against 49.9% of low-confidence canonicals. Emergence-cohort canonicals are enriched for established homomers, 75.0% against 48.7% for other canonicals of the same confidence, so emergence marks full-length predictions that fail and not interfaces created by truncation. Truncation-associated remodeling of homodimer architecture is a graded, topology-asymmetric property of the human proteome, readable from interface size and operating across gene-functional categories.

## Introduction

Most human protein-coding genes produce multiple isoforms through alternative splicing, alternative promoter usage and post-translational proteolysis, resulting in a proteome of tens of thousands of distinct protein products whose individual structural and functional consequences remain only partially mapped (Wang et al. 2008; Pan et al. 2008; Yang et al. 2016; Liu et al. 2022; Higdon et al. 2024). Among these consequences, changes in the oligomeric organization at which proteins assemble for catalysis, allostery, signal transduction and chromatin engagement, are particularly consequential because they reorganize function rather than merely modulating it (Buljan et al. 2012; Marsh and Teichmann 2015). Whether and how truncation alters quaternary state has, however, been examined largely on a molecule-by-molecule basis, and what has been missing is a systematic, statistically grounded framework that places these individual cases within a proteome-wide grammar.

Tools like AlphaFold-Multimer enable structural prediction of user-defined oligomers from sequence alone (Jumper et al. 2021; Evans et al. 2022; Abramson et al. 2024). Proteome-scale prediction of homo-oligomers has been performed before, including an atlas spanning four proteomes of which two are eukaryotic (Schweke et al. 2024), and a recent expansion of the AlphaFold Protein Structure Database performed in collaboration with NVIDIA now populates predicted homodimer structures across the human proteome, differing in scale and, for the present purpose, in covering annotated isoform accessions alongside canonical ones (Varadi et al. 2024; Han et al. 2026). These resources allow, for the first time, a one-to-one comparison of every annotated truncated isoform against its canonical homodimer counterpart in a single common-confidence framework, and remove the central historical obstacle to systematic isoform-level structural comparison.

Structural case studies in individual receptor and transporter families have indicated that truncated isoforms can retain key features of the canonical fold (Qing et al. 2020) and can alter or inhibit the oligomeric assembly of their canonical counterparts (Li et al. 2023; Li et al. 2024; Karagöl, A. et al. 2024; Karagöl, T. & Karagöl, A. 2026c), and can depart from the canonical fold to assemble into a state of their own (Karagöl, A. & Karagöl, T. 2025a). Because each of these studies was confined to a single gene family, whether truncation-driven changes in quaternary state follow general rules has remained open, and the present work addresses this question across the entire predicted human homodimer landscape.

Truncation here means an annotated shorter protein product, generated by alternative splicing, an alternative promoter or post-translational cleavage, and not a nonsense or frameshift variant in the sense used in clinical genetics. We scored 5,168 canonical-versus-truncated homodimer pairs across the human proteome on each of the four interface metrics the release distributes. We asked whether the truncation landscape partitions into distinct state-transition classes, and whether any such partition survives on predicted interface size instead of resting on a confidence score alone. We evaluated topological asymmetry between N-terminal and C-terminal sequence losses, and tested whether it is independent of how much sequence a deletion removes. We quantified which curated structural features the deletions remove, against a null in which windows of the same length fall at random positions in the same protein. We mapped the resulting classes onto biological networks to establish whether the rules operate broadly across functional categories or concentrate within particular pathways.

Placing truncation on a proteome-scale structural footing has broad implications. The framework supplies a statistical prior for unannotated truncated isoforms, in which deletion position and deletion magnitude together predict whether a homodimer interface survives, applicable to prioritizing targets in transcriptomic atlases, cancer splicing landscapes and disease-associated alternative-splicing programs. The same framework offers a principled basis for rational protein engineering alongside complementary approaches such as computational epistatic rescue of destabilizing substitutions and reverse-QTY conversion of a soluble fold into a membrane-compatible one (Karagöl, A. & Karagöl, T. 2026c; Karagöl, T. & Karagöl, A. 2026a). In disease biology, truncation-associated ablation and emergence events form a structurally articulated reservoir of splicing-driven phenotypes amenable to systematic prediction in oncology, neurodegeneration and immune-system biology, where alternative-splicing programs generate disease-associated isoform profiles. For therapeutic development, the subset of emergence events that reproduce across independent interface metrics defines a tractable set of sequence-encoded, conditionally exposed homodimer interfaces open to targeted pharmacological modulation. This structural grammar supplies a statistical reference frame for subsequent computational and experimental work on the transitions most consequential to human biology and medicine.

## Results and Discussions

### Proteome-wide canonical and truncated homodimer landscape

We constructed a canonical-versus-truncated homodimer analysis by pairing every AlphaFold-Multimer scored canonical homodimer with each annotated truncated isoform of the same gene, scoring each truncated isoform under identical analysis parameters as a homodimer constructed from two copies of the truncated sequence. This produced 5,168 (canonical, isoform) pairs across the human proteome, to our knowledge among the largest one-to-one canonical-versus-truncated homodimer comparisons reported to date and substantially larger than any previously published focused-family analysis. To stratify outcomes without resorting to a binary "dimer or not" classifier, we partitioned canonical homodimers into three confidence tiers based on canonical interface predicted Template Modeling score (ipTM): a High-confidence tier (n = 234; canonical ipTM ≥ 0.8), a Moderate tier (n = 354; 0.6 ≤ canonical ipTM < 0.8), and a Low tier (n = 4,580; canonical ipTM < 0.6). Each (canonical, isoform) pair was then assigned a state-transition class based on the combined tier of its two endpoints (Figure 1). Pairing was performed strictly within each base-accession family of UniProt accessions (the primary accession after stripping any trailing isoform suffix), so that both members of every pair share locus assignment by construction (Table S1).

**Figure 1.**
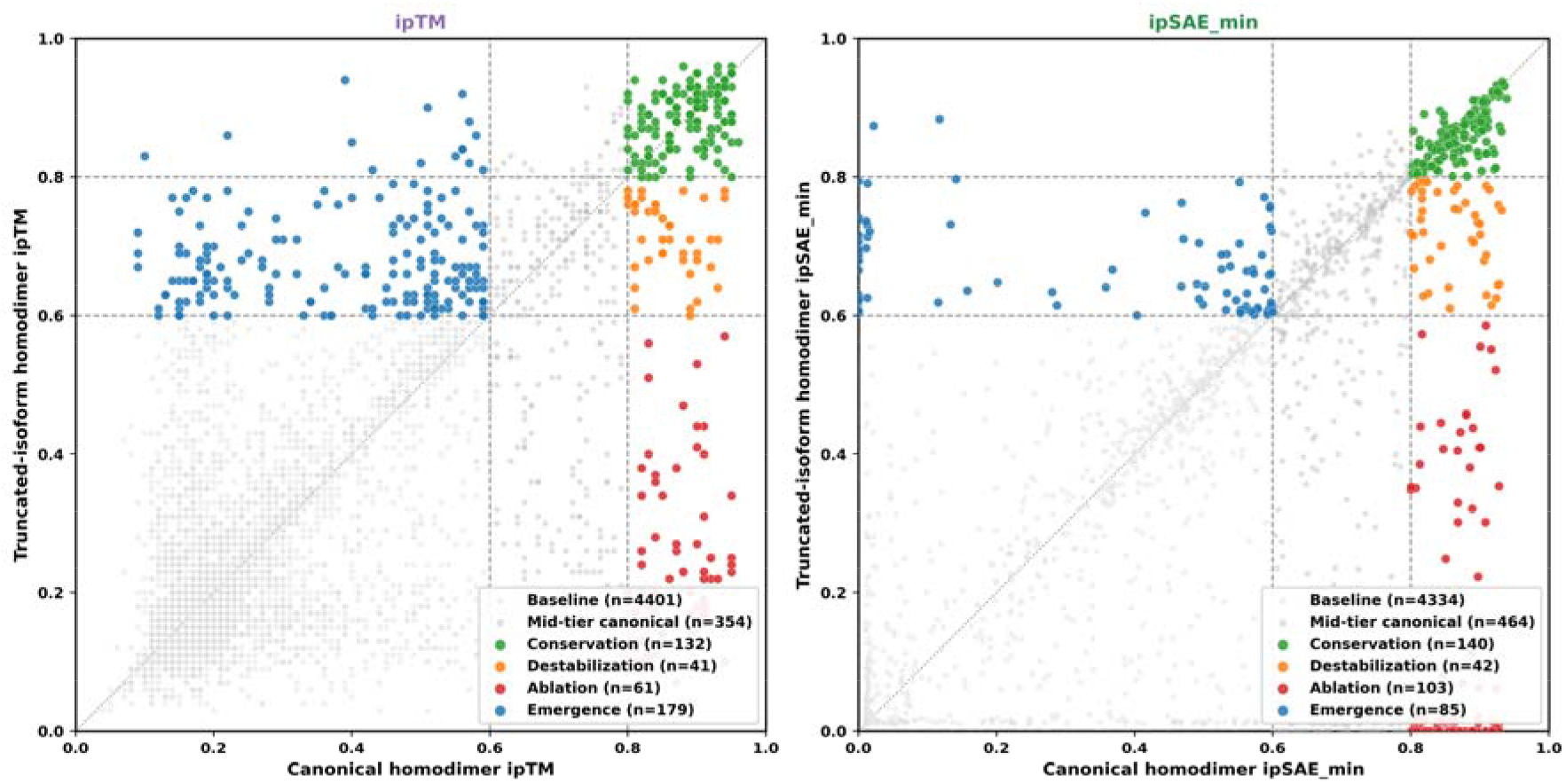
Canonical-versus-isoform landscape on two interface metrics. Canonical homodimer score (x-axis) against truncated-isoform homodimer score (y-axis) across all 5,168 (canonical, isoform) pairs. Left, ipTM. Right, ipSAE, the metric on which the release defines its own confidence tiers. Each point is one pair, colored by state-transition class under identical rules on each metric, with class sizes given in the legend. Emergence here counts the 179 pairs rising from a sub-threshold canonical, and the 208-event cohort used for the annotation analyses adds the 29 tier-promotion events. Applying the same tier rules to ipSAE leaves 85 in that class, while 42 of the 208 reproduce at the release threshold (Figure 4). Dashed reference lines mark the 0.6 and 0.8 thresholds on each axis, and the dotted diagonal marks no change. The emergence class contracts and the complete-ablation class expands when the same rules are applied to ipSAE.

### Four interface metrics and what each one supports

The release distributes four interface quantities for every predicted complex: ipTM, ipSAE, pDockQ and LIS, together with counts of inter-chain contacts and of residues inside the interface distance cutoff. It defines its own high-confidence set and its own confidence tiers on ipSAE, which corrects a known dependence of ipTM on chain length and on poorly predicted segments (Dunbrack 2025). We therefore report every classification on ipTM and on ipSAE in parallel, and use the interface residue counts as an independent structural quantity that no confidence model enters.

The four metrics do not move together equally. Across all 5,168 pairs the change in ipSAE on truncation correlates with the change in LIS at Spearman rho = 0.840 and with the change in ipTM at 0.654, while pDockQ correlates weakly with all three (0.284 to 0.343). Thresholds cannot be transferred between metrics on different scales, so we also classified each pair under thresholds matched by quantile to the ipTM tiers. Under that matching the conservation class holds 137 pairs on ipTM and 110 on ipSAE, the complete-ablation class 71 and 90, and the emergence class 184 and 67.

### Classification of homodimer state transitions

Stratifying the 5,168 pairs by canonical tier produced four reproducible state-transition classes (Table 1). In the 234 High-confidence canonical homodimers, truncation was associated with structural conservation in 132 pairs (56.4%), predicted complete interface ablation in 61 (26.1%), and partial destabilization to Moderate confidence in 41 (17.5%) (Tables S3, S4), three classes that together account for the entire High-tier population. Of the 132 conservation events, 100 remained conserved when the same rule was applied to ipSAE. The three-class partition replicated qualitatively in the 354 Moderate-confidence canonicals. In the 4,580 Low-confidence canonicals the dominant outcome was non-transition, with a smaller branch in which a sub-threshold canonical homodimer became a predicted Moderate-confidence or High-confidence homodimer after truncation. 179 pairs meet that definition, and we refer to them as truncation-associated emergence. 29 further pairs had canonical ipTM between 0.60 and 0.79 and were tier promotions inside the confident range; they were reported with the conservation class and were excluded from the emergence analyses below.

**Table 1.** Four-class state-transition assignment of (canonical, isoform) homodimer pairs.

| Canonical tier | Tier transition | Outcome class | Pairs (n) | % of canonical-tier total | Mean ΔipTM | Mean residues lost |
|---|---|---|---|---|---|---|
| High (n = 234) | High → High | Conservation (canonical preserved) | 132 | 56.4% | +0.000 | 85.0 |
|  | High → Moderate | Partial destabilization | 41 | 17.5% | −0.147 | 133.9 |
|  | High → Low | Complete interface ablation | 61 | 26.1% | −0.618 | 177.6 |
| Moderate (n = 354) | Moderate → Moderate | Conservation (Moderate preserved) | 158 | 44.6% | +0.008 | 86.8 |
|  | Moderate → High | Strict emergence (gain) | 29 | 8.2% | +0.107 | 149.7 |
|  | Moderate → Low | Ablation from Moderate | 167 | 47.2% | −0.380 | 187.5 |
| Low (n = 4,580) | Low → Low | Non-dimer baseline | 4,401 | 96.1% | −0.005 | 157.4 |
|  | Low → Moderate | Strict emergence (gain) | 164 | 3.6% | +0.283 | 248.9 |
|  | Low → High | Strict emergence (gain) | 15 | 0.3% | +0.381 | 276.9 |

Stratification of all 5,168 (canonical, isoform) pairs by canonical ipTM tier (High ≥ 0.8; Moderate 0.6-0.8; Low < 0.6) and ending-tier outcome. The four reproducible state-transition classes, Conservation, Partial destabilization, Complete ablation, and Strict emergence (gain), are highlighted by row shading (green = preserved/gained, amber = partially destabilized, red = ablated). Tier-conditional percentages are computed within each canonical-tier total. Mean ΔipTM is isoform ipTM minus canonical ipTM; positive values are conserved/gained, negative values indicate loss. Residues lost are alignment-derived (canonical length minus isoform length, summed within class).

The relationship between deletion magnitude and predicted interface fate was graded, and it was graded on interface size and not only on interface confidence (Figure 2). Across the 749 pairs whose canonical has a detectable interface by ipSAE, the fraction of canonical interface residues retained by the isoform falls monotonically with the fraction of the chain removed: 0.997 below a tenth, 0.883 between a tenth and a fifth, 0.780 between a fifth and three tenths, 0.489 between three tenths and a half, and 0.192 above a half (Spearman rho = -0.552, p = 4.5 × 10^−61^). The three High-tier outcome classes separate on the same quantity, with conservation events retaining a median 0.938 of the canonical interface, destabilization events 0.622, and ablation events 0.000 (Kruskal-Wallis p = 5.8 × 10^−32^). Interface confidence tracks interface extent across these pairs, at Spearman rho = 0.658 for ipTM and 0.700 for ipSAE. Truncation magnitude was therefore a graded determinant of predicted interface fate, and the grading was visible in the predicted contacts themselves. The gradient was not an artefact of chain length. Across the 2,184 pairs whose canonical has a detectable interface, the fraction of the chain removed correlates with the change in ipTM at Spearman rho = -0.155, and with the change in interface residue count at -0.351. Holding the change in interface residues constant removes the first correlation (partial rho = +0.094), so the ipTM gradient with deletion size was driven by interface loss and not by the length term in the score.

**Figure 2.**
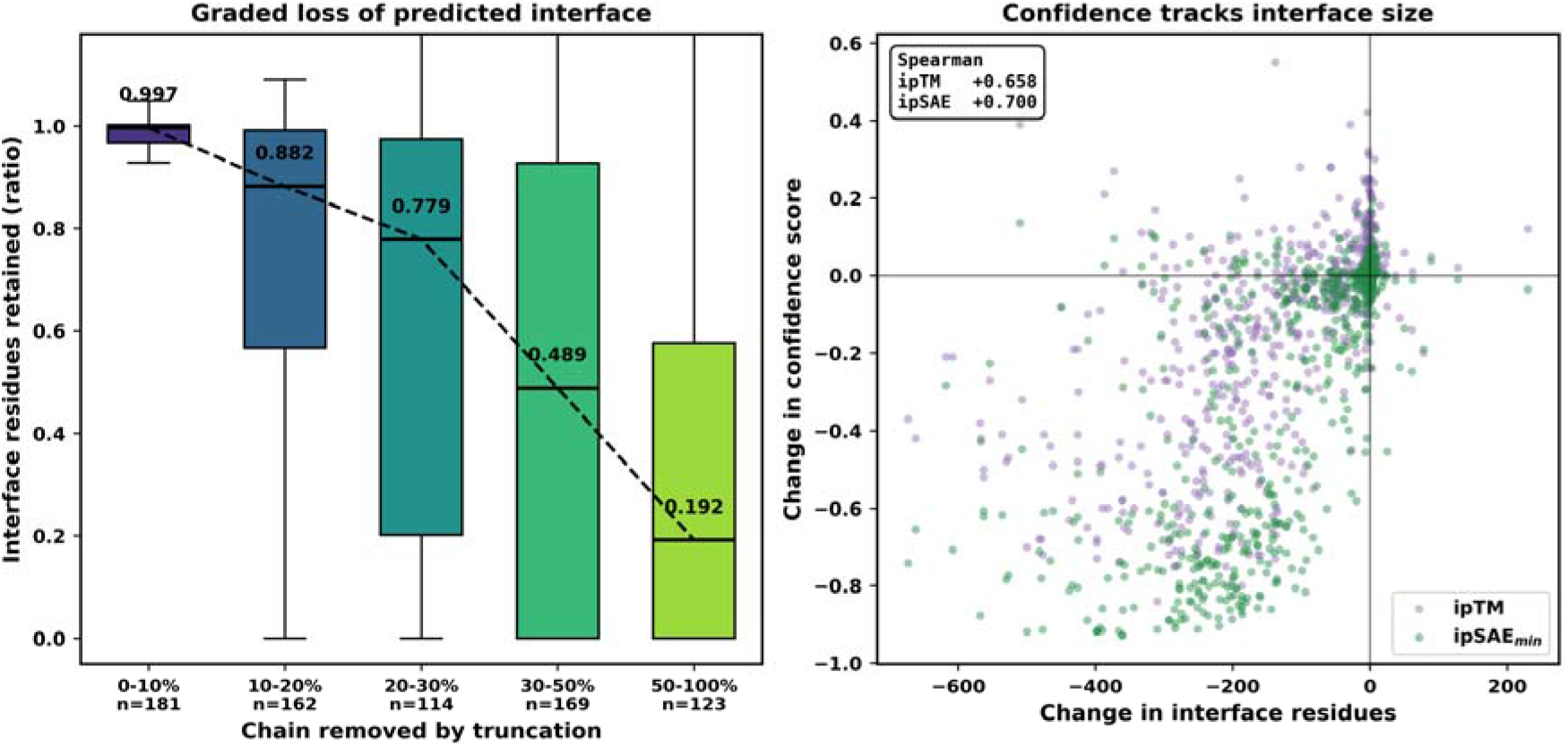
Graded loss of predicted interface with truncation magnitude. (a) Fraction of canonical interface residues retained by the isoform, binned by the fraction of the chain the deletion removes, across the 749 pairs whose canonical has a detectable interface by ipSAE. Medians are printed and joined. (b) Change in interface confidence against change in the count of interface residues, for ipTM and for ipSAE, with Spearman coefficients inset.

First, the consistent replication of outcome distributions across both High and Moderate canonical-tier baselines indicates that this behavior is likely intrinsic to the truncation events themselves. Rather than reflecting the idiosyncrasies of a specific protein family or fold class, these predictions suggest the emergence of a generalized structural grammar governing homodimer assembly.

Second, the observed continuous dose-response relationship fundamentally reframes how truncation might be conceptualized at the structural level. In genetics and transcriptomics, truncation-bearing splice isoforms are typically treated as binary perturbations, with an implicit assumption that their structural consequences act as discrete switches. However, at the level of predicted homodimer architecture, truncation appears to behave instead as a quantitative lever spanning a graded landscape. In this proposed model, truncations frequently preserve homodimeric capacity, acting as a structural rheostat in which the magnitude of a deletion sets the expected loss of interface and its position modulates that loss. Within this landscape, a distinct trajectory suggests the formation of homodimeric interfaces not confidently predicted in the canonical form. These predictions map the landscape of homodimer architectures and make no implications regarding inhibitory functions. Because this framework models truncated isoforms in isolation to evaluate their independent homodimeric potential, future studies will be required to explore their potential interactions with canonical counterparts, which case studies in individual transporters suggest can be inhibitory or competitive (Karagöl, A. & Karagöl, T. 2025b, 2026a), and to determine whether these variants participate in heterodimer networks. The natural conceptual model at a proteome scale therefore shifts from knockout-by-splicing to rheostat-by-splicing, and the partition reproduces across both High and Moderate canonical tiers instead of being confined to one.

### Agreement between predicted confidence and deposited structures

Where deposited structures exist, they agree with the predicted confidence. Of the 4,115 canonical accessions, we found 2,058 with at least one entry in the Protein Data Bank (Berman et al. 2000), spanning 22,277 entries, and deposited chains were mapped to accessions through SIFTS (Dana et al. 2019). For each accession we took the largest number of copies of that protein in any deposited biological assembly. Among canonicals with a structure, two or more copies appear for 95.3% of the High-confidence tier (n = 127), 85.4% of the Moderate tier (n = 164) and 49.9% of the Low tier (n = 1,767) (Fisher exact p = 1.5 × 10^−27^; Table S13). Restricting to assemblies composed only of that protein gives 92.1%, 76.8% and 36.2%. Splitting instead on ipSAE_min at 0.6 gives 84.8% against 49.6% (p = 3.7 × 10^−36^). Two or more copies in a deposited assembly is a necessary condition for a homodimer and not a sufficient one, since crystal packing can place copies in contact, and structural coverage is uneven because proteins that crystallize readily are over-represented. The comparison bounds agreement between the predicted tiers and deposited assemblies, and does not establish the oligomeric state of any individual protein.

### Topological determinants of homodimer preservation

Re-deriving every deletion window by global alignment, the 5,164 pairs with both sequences available partition into 2,750 N-terminal losses (53.3%), 2,207 C-terminal losses (42.7%), 112 internal or splice deletions (2.2%) and 95 events removing sequence at both termini (1.8%) (Table S2; Figure 3). The C-terminal share was stable between 42.7% and 43.3% as the terminal tolerance was swept from 0 to 20 residues, and rose to 46.1% when every pair whose deletion coincided with an annotated cleavage site was excluded. Exact prefix and suffix matching assigns a far smaller C-terminal share, because splicing commonly leaves a few novel junction residues before the stop codon, and a window that stops short of the final residue is then read as an internal deletion.

**Figure 3.**
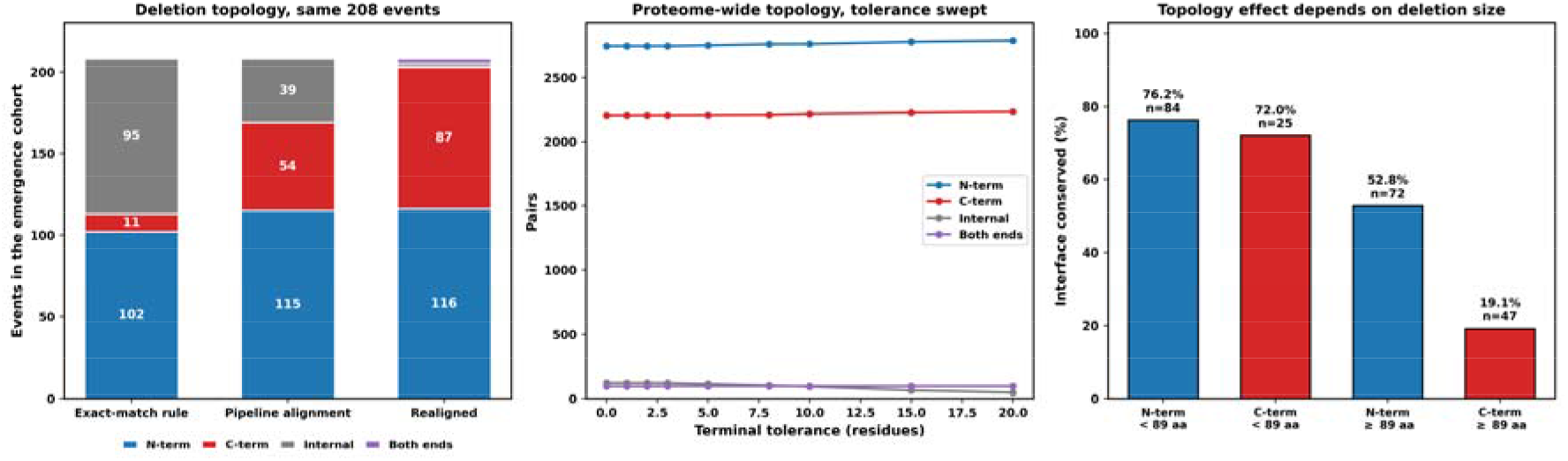
Deletion topology, its stability, and its effect on interface conservation. (a) Topology of the same 208 emergence events under three assignments: the exact-match rule, the alignment used in the mapping pipeline, and realignment by affine-gap global alignment with free end gaps. C-terminal events are under-called by both of the first two, because a deletion that leaves novel junction residues before the stop codon produces a window stopping short of the final residue. (b) Proteome-wide topology across all 5,164 realigned pairs as the terminal tolerance is swept from 0 to 20 residues; the C-terminal share moves between 42.7% and 43.3%. (c) Interface conservation rate by topology among canonicals at ipTM 0.8 and above, split at the cohort median deletion of 89 residues. The topology asymmetry appears only for the larger deletions.

Topology predicts outcome class. Among the 234 High-confidence canonicals, interface conservation follows 65.4% of N-terminal losses (n = 156) and 37.5% of C-terminal losses (n = 72), a bootstrap 95% interval of 57.7 to 72.4 versus 26.4 to 48.6 (Figure 3). The asymmetry depends on how much sequence the deletion removes. For deletions below the cohort median of 89 residues, conservation follows 76.2% of N-terminal and 72.0% of C-terminal losses. At or above that median the two diverge, to 52.8% and 19.1%. Position therefore mattered most where the deletion was large, and small deletions at either terminus were comparably well tolerated.

The destabilization and ablation signatures were most visible in classical homodimeric metabolic enzymes, and the deletions that produce them reach a chain terminus (Table S4; Table 2). The UDP-glucuronosyltransferase locus contributes nine events across the two classes, under the symbols UGT1A1, UGT1A3, UGT1A4, UGT1A5, UGT1A6, UGT1A7, UGT1A8, UGT1A9 and UGT1A10. Each removes the same 89 residues and each drops the predicted dimer ipTM by 0.19 to 0.29. These paralogues share the terminal exons of one locus, so the nine events report a single splicing event, and grouping by the identity of the deleted segment collapses them to one (Table S12). Every one of the nine deletions runs to the final canonical residue with eight substituted residues at the junction, which an exact suffix test reads as an internal deletion. Other destabilization leads follow both terminal modes: FMO5 falls from 0.93 to 0.71 after a 252-residue C-terminal loss and EPM2A from 0.90 to 0.68 after a 14-residue C-terminal loss, alongside the N-terminal losses of CRYZ (0.94 to 0.77 after 137 residues), ASRGL1 (0.94 to 0.78 after 128), GSTA4 (0.93 to 0.64 after 93), AOC3 (0.92 to 0.71 after 543) and FAP (0.91 to 0.77 after 521). HEXA (0.92 to 0.67) and one RHBG product remove sequence at both termini. Many of these enzymes adopt TIM-barrel or Rossmann-fold architectures in which interface integrity depends on extended contact surfaces across the dyad axis (Ruan et al. 2015; Liu and Tanner 2019; Shortall et al. 2021), so a deletion reaching either terminus can compromise core interface contacts.

**Table 2.** Top 15 partial-destabilization leads from the High canonical-tier baseline.

| Gene | Canonical UniProt | Isoform UniProt | Canonical ipTM | Isoform ipTM | ΔipTM | Residues lost | Topology | Outcome |
|---|---|---|---|---|---|---|---|---|
| **CRYZ** | Q08257 | Q08257-2 | 0.94 | 0.77 | −0.17 | 137 | N-terminal loss | Destabilization |
| **ASRGL1** | Q7L266 | Q7L266-2 | 0.94 | 0.78 | −0.16 | 128 | N-terminal loss | Destabilization |
| **FMO5** | P49326 | P49326-2 | 0.93 | 0.71 | −0.22 | 248 | C-terminal loss | Destabilization |
| **GSTA4** | O15217 | O15217-2 | 0.93 | 0.64 | −0.29 | 93 | N-terminal loss | Destabilization |
| **AOC3** | Q16853 | Q16853-3 | 0.92 | 0.71 | −0.21 | 543 | N-terminal loss | Destabilization |
| **HEXA** | P06865 | P06865-2 | 0.92 | 0.67 | −0.25 | 361 | Both termini | Destabilization |
| **FAP** | Q12884 | Q12884-2 | 0.91 | 0.77 | −0.14 | 521 | N-terminal loss | Destabilization |
| **UGT1A7** | Q9HAW7 | Q9HAW7-2 | 0.90 | 0.62 | −0.28 | 89 | C-terminal loss | Destabilization |
| **UGT1A9** | O60656 | O60656-2 | 0.90 | 0.69 | −0.21 | 89 | C-terminal loss | Destabilization |
| **EPM2A** | O95278 | O95278-2 | 0.90 | 0.68 | −0.22 | 14 | C-terminal loss | Destabilization |
| **RHBG** | Q9H310 | Q9H310-4 | 0.89 | 0.71 | −0.18 | 69 | N-terminal loss | Destabilization |
| **RHBG** | Q9H310 | Q9H310-5 | 0.89 | 0.61 | −0.28 | 156 | Both termini | Destabilization |
| **UGT1A1** | P22309 | P22309-2 | 0.89 | 0.66 | −0.23 | 89 | C-terminal loss | Destabilization |
| **UGT1A3** | P35503 | P35503-3 | 0.89 | 0.60 | −0.29 | 89 | C-terminal loss | Destabilization |
| **UGT1A4** | P22310 | P22310-2 | 0.88 | 0.69 | −0.19 | 89 | C-terminal loss | Destabilization |

Top 15 (canonical, isoform) pairs from the 41-event High-tier partial-destabilization class (High → Moderate), ranked by canonical ipTM. The list is dominated by classical homodimeric metabolic enzymes whose dimer interface is destabilized predominantly by internal-splice deletions (the UGT1A glucuronosyltransferase family with recurrent ~89-residue splice losses, plus CRYZ, FMO5, HEXA, EPM2A, RHBG) or by N-terminal losses (ASRGL1, GSTA4, AOC3, FAP, RHBG), consistent with the destabilization signature reported in Results. Full ranked list of 41 events is provided as Supplementary Table.

This observed asymmetry is consistent with well-documented properties of human proteins at the case level. N-terminal signal peptides, propeptides, transactivation domains and disordered low-complexity segments are common in eukaryotic proteomes and are, in many specific cases, structurally dispensable for the integrity of canonical folds. Conversely, several major enzyme architectures, including TIM-barrel and Rossmann-fold homodimers, depend on extended interface contacts across the dyad axis whose disruption by internal-splice deletions or by removal of structurally extended N-terminal segments is well documented, and our destabilization-leads cohort (Table 2) is dominated by enzymes of these classes. We emphasize that the N-terminal-preference for conservation is a tendency rather than a universal rule of protein architecture: well-documented counterexamples exist, including N-terminal coiled-coil dimerization motifs in transcription factors and viral fusion proteins (Blobel and Dobberstein 1975; Ellenberger et al. 1992; Chan et al. 1997), and C-terminal regulatory tails in Src-family kinases (Sicheri et al. 1997), RNA polymerase II (Cramer et al. 2001) and a range of nuclear hormone receptors (Brzozowski et al. 1997). What our data establish is that the N-terminal-dispensability tendency holds on average, across 5,168 predicted human homodimers, and at sufficient resolution to support its use as a structural prior. C-terminal events are not rare, at 42.7% of the aligned pairs, and the asymmetry they carry is in outcome and not in frequency: interface conservation follows 65.4% of N-terminal losses against 37.5% of C-terminal losses, and only for deletions above the cohort median of 89 residues.

Because this structural organization is widely conserved, truncation topology naturally emerges as an informative prior for modeling homodimer assembly. For an unannotated truncated isoform evaluated in isolation, the geometry of the deletion alone provides a statistical basis to anticipate its independent structural fate. Isoforms experiencing N-terminal loss are predisposed to retain or gain homodimeric architecture, whereas those with C-terminal loss are predisposed toward homodimer interface attenuation. Crucially, these predictive patterns map exclusively to independent homodimer formation; they do not address potential heterodimerization with canonical sequences or inhibitory dynamics, which remain distinct avenues for future investigation. This proteome-wide quantification places the family-level case studies cited in the Introduction on a statistical footing. While specific structural exceptions naturally exist across the dataset, the macroscopic pattern remains highly reproducible, offering a reliable heuristic for the predictive modeling of isolated structural splice variants.

### Truncation-driven emergence cohort

Truncation-associated emergence covered 179 pairs in which a sub-threshold canonical homodimer was predicted with Moderate or High confidence after truncation, together with the 29 tier-promotion events described above, and the 208-event union is the population used for the annotation analyses that follow (Table S5, Table 3). The cohort spans 188 distinct gene products. Grouping events by the identity of the deleted segment finds no paralogous duplication within it at any identity threshold between 0.80 and 0.99, in contrast to the destabilization and ablation classes, where the UGT1A locus contributes one splicing event under six and three gene symbols respectively.

**Table 3.** Top 15 strict-emergence (truncation-induced gain) leads.

| Gene | Canonical UniProt | Isoform UniProt | Canonical ipTM | Isoform ipTM | ipTM gain | Residues lost | Topology | Tier transition |
|---|---|---|---|---|---|---|---|---|
| **THAP4** | Q8WY91 | Q8WY91-2 | 0.39 | 0.94 | +0.55 | 412 | N-terminal loss | Low → High |
| **RIOX1** | Q9H6W3 | Q9H6W3-2 | 0.74 | 0.93 | +0.19 | 212 | N-terminal loss | Moderate → High |
| **CBR4** | Q8N4T8 | Q8N4T8-2 | 0.56 | 0.92 | +0.36 | 58 | C-terminal loss | Low → High |
| **B4GALNT2** | Q8NHY0 | Q8NHY0-3 | 0.79 | 0.90 | +0.11 | 86 | N-terminal loss | Moderate → High |
| **PDE8B** | O95263 | O95263-5 | 0.51 | 0.90 | +0.39 | 535 | N-terminal loss | Low → High |
| **PRDX5** | P30044 | P30044-2 | 0.74 | 0.90 | +0.16 | 52 | N-terminal loss | Moderate → High |
| **NUDT6** | P53370 | P53370-2 | 0.79 | 0.89 | +0.10 | 169 | N-terminal loss | Moderate → High |
| **CAPN3** | P20807 | P20807-5 | 0.78 | 0.89 | +0.11 | 665 | N-terminal loss | Moderate → High |
| **BDH2** | Q9BUT1 | Q9BUT1-3 | 0.78 | 0.88 | +0.10 | 101 | N-terminal loss | Moderate → High |
| **GFER** | P55789 | P55789-2 | 0.57 | 0.88 | +0.31 | 80 | N-terminal loss | Low → High |
| **TFR2** | Q9UP52 | Q9UP52-2 | 0.78 | 0.88 | +0.10 | 171 | N-terminal loss | Moderate → High |
| **ACOT12** | Q8WYK0 | Q8WYK0-2 | 0.22 | 0.86 | +0.64 | 388 | C-terminal loss | Low → High |
| **FH** | P07954 | P07954-2 | 0.58 | 0.86 | +0.28 | 43 | N-terminal loss | Low → High |
| **ANO3** | Q9BYT9 | Q9BYT9-2 | 0.77 | 0.85 | +0.08 | 146 | N-terminal loss | Moderate → High |
| **HAAO** | P46952 | P46952-2 | 0.40 | 0.85 | +0.45 | 139 | C-terminal loss | Low → High |

Top 15 (canonical, isoform) pairs from the 208-event strict-emergence class (Low/Moderate → Moderate/High), ranked by isoform ipTM. The cohort is dominated by N-terminal-loss isoforms (11/15 here; 49.0% of the full 208-event class versus 46.6% baseline), with Low → High events (THAP4, ACOT12) showing the largest ipTM gains. Full 208-event cohort is provided as Supplementary Table.

Whether the class reflects a structural transition or the behavior of one confidence model is answerable from the release itself. Applying the same rule to ipSAE, the metric on which the release defines its tiers, 42 of the 208 events reproduce (Figure 4). Those 42 separated distinctly from the remainder on predicted interface extent: their median gain was 25 interface residues, against zero for the other 166, and their median ipSAE gain was 0.504 versus 0.047. Direction of change was more widely shared than threshold crossing, with 115 of the 208 events increasing on all four metrics and 178 on at least three (Table S10). The events that reproduce split 22 N-terminal and 20 C-terminal, so emergence was not specific to loss of N-terminal sequence. Twenty-five of the 208 deleted an annotated cleavage site, 11 a signal peptide, 11 a transit peptide and 7 a propeptide; PRDX5 and FH were of this kind, where the isoform UniProt names as cytoplasmic is the mature chain after removal of a mitochondrial targeting sequence. Reproduction tracks how far the isoform clears the threshold: 27 events whose isoform ipTM lies between 0.60 and 0.62 yield 4 that reproduce (14.8%), against 22 of the 85 that exceed 0.70 (25.9%). Events whose deletion coincides with an annotated cleavage site behave like the rest of the cohort: 7 of 25 reproduce on ipSAE (28.0%) against 35 of 183 (19.1%), with no detectable difference between them, so removal of a targeting peptide does not account for the reproducing subset.

**Figure 4.**
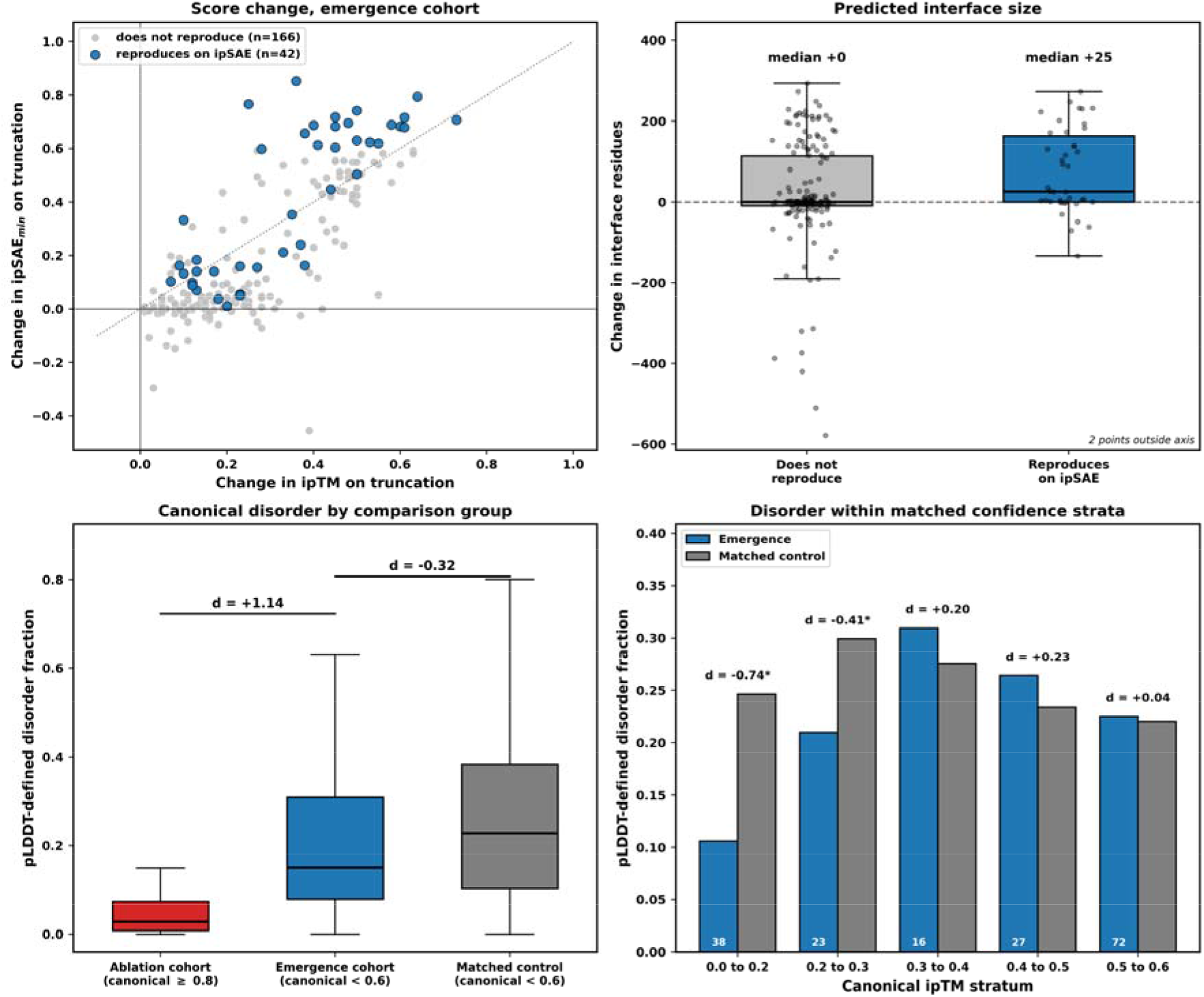
Which emergence events reproduce on the release’s own metric. (a) Change in ipSAE against change in ipTM for the 208 emergence events, with the 42 that reproduce on ipSAE at the release threshold highlighted. (b) Change in the count of predicted interface residues for the events that reproduce against those that do not; medians are marked and points outside the axis are noted. (c) Canonical pLDDT-defined disorder fraction for the complete-ablation cohort, the emergence cohort, and controls drawn from the same sub-threshold canonical stratum as the emergence cohort, with Cohen’s d for each contrast. (d) The same comparison within narrow canonical ipTM strata, with Cohen’s d annotated and asterisks marking p below 0.05; query sizes are printed inside the bars.

The emergence cohort was enriched for proteins that deposited structures already show to be homomers. Of the 188 canonical gene products in the cohort, we found structures for 112, and 84 of those (75.0%) appeared in two or more copies in a deposited assembly, against 816 of 1,674 (48.7%) among other canonicals below the same confidence boundary (Fisher exact p = 4.8 × 10 ). The difference held within every stratum of structural coverage, with the emergence rate running from 50.0% among proteins carrying a single entry to 88.2% among those carrying ten or more, so it was not an artefact of how much structural data each protein has. On these proteins the truncated-isoform prediction agrees with the deposited assembly and the full-length prediction does not. Events that reproduce on ipSAE were no different in this respect from those that did not, at 13 of 18 (72.2%) against 72 of 95 (75.8%) (Fisher exact p = 0.769). The cohort was selected on the isoform reaching the threshold, so part of the enrichment follows from that selection, and the comparison does not show truncation creating an interface where none exists. The reproducing events whose canonical is monomeric in every deposited assembly, ALCAM, CHIA, EPHB4, HAAO and HLA-G, were the subset for which a truncation-dependent interface remains open.

### Biophysical divergence between emergence and ablation

Canonical disorder content was computed as the fraction of residues with AlphaFold pLDDT below 50, read from the AFDB prediction metadata for all 4,115 canonical accessions (Table S6). Comparing the emergence cohort against the high-confidence ablation cohort reproduces a large separation, 0.1997 versus 0.0437 with Cohen’s d = 1.14 (p = 5.0 × 10 ¹ ). The two cohorts were drawn from opposite ends of the canonical confidence range by construction, and disorder depresses interface confidence, so that contrast was not independent of the selection. Against 4,324 controls drawn from the same sub-threshold canonical stratum as the emergence cohort, the difference was small and ran the other way, 0.1997 versus 0.2613 with Cohen’s d = -0.32 (p = 2.2 × 10 ). Within narrow canonical ipTM strata there was no detectable difference above 0.3, with Cohen’s d between +0.04 and +0.23, and in the two lowest strata the emergence canonicals showed less disorder (d = -0.74 and -0.41). Canonical disorder also shows no detectable difference between the 42 events that reproduce on ipSAE and the 166 that do not (d = -0.17, p = 0.093). Across the whole dataset, disorder and canonical ipTM were weakly anticorrelated (Spearman rho = -0.101, p = 5.4 × 10 ¹³), which is enough to generate a large apparent separation between cohorts selected at opposite ends of that range.

Peak per-residue binding propensity, taken as the single highest score anywhere in a sequence, scaled with sequence length across the profiled cohorts (Spearman rho = 0.349, p = 3.9 × 10 ), and the emergence isoforms were the longer group, averaging 337 residues versus 263 for the ablation isoforms. Neither disorder measure therefore separates the emergence class from a properly matched comparison group. What distinguished the events that reproduce across metrics was the predicted interface itself, which grew in those events and was unchanged in the rest.

### Predominance of disorder and compositional bias in deleted segments

To resolve which annotated structural features truncation removes, we intersected every alignment-derived deletion window in the emergence cohort with the canonical UniProt feature set across the retained feature classes (Table S7). 88.5% of events delete at least one curated feature, 53.4% delete a disordered or compositional-bias region, and 1.0% delete a feature annotated as inhibitory, regulatory or pseudo-substrate (Figure S1; Table S11). Deletions in this cohort were large, removing on average a substantial share of the chain, so the rate at which they strike curated features must be read against the rate a window of the same length would achieve at a random position in the same protein. Drawing 2,000 such windows per event gives 86.0% for any curated feature, 42.7% for disorder or compositional bias, and 1.4% for the inhibitory class. The overall feature-deletion rate was therefore close to what window size alone produced. Disordered and compositional-bias regions were struck about ten points more often than chance, and annotated inhibitory modules showed no excess.

The dominant deleted feature classes, ranked by frequency of intersection events, were Region (247 instances), Compositional bias (200), Binding site (160), Domain (108), Topological domain (81), Transmembrane (62), Repeat (50), Motif (30), Active site (27), Coiled coil (27), Zinc finger (18), Propeptide (7) and DNA binding (5). Deletions overlapped a disordered or compositional-bias region in 53.4% of events, where a window of the same length placed at random within the same canonical sequence gave 42.7%, while the inhibitory, regulatory and pseudo-substrate class was deleted in 1.0% of events where the same null gave 1.4% and showed no excess over that null. The rate did not depend on where the deletion fell: disordered or compositional-bias sequence was removed in 50.0% of N-terminal events (n = 116) and 57.5% of C-terminal events (n = 87), with no detectable difference between them (Fisher exact p = 0.321). The disorder signal and the topology asymmetry were therefore separate effects and not one effect read twice. The conventional auto-inhibitory domain release model, in which truncation activates a latent function by removing a curated inhibitory module, therefore accounts for about 1.0% of events in this cohort. What these deletions remove instead is disordered and compositional-bias sequence, which UniProt rarely curates as inhibitory. One reading consistent with this pattern is that a latent interface is occluded by a disordered or compositional-bias segment whose removal exposes it, which would be auto-inhibitory in a structural and not an annotation-based sense. These data locate the deleted sequence and its annotation class and do not resolve the interface before and after the deletion, so occlusion remains a hypothesis they motivate and do not settle.

Four events at the connectivity core of the cohort differ in whether the rise in ipTM reproduces on the length-corrected metric. EPHB4 loses 573 residues, almost all of them in a single block running from residue 419 to the C terminus, which removes the transmembrane helix at residues 540 to 560 along with the entire cytoplasmic region with the kinase and SAM domains and leaves a truncated extracellular segment. The soluble EphB4 ectodomain of this kind antagonizes EphB4-ephrinB2 interaction and modulates angiogenesis (Kertesz et al. 2006), so the truncated form corresponds to a characterized protein product. Its predicted interface rises on both metrics, from 0.20 to 0.70 on ipTM and from 0.00 to 0.630 on ipSAE, crossing the release threshold on each. EPHA2 loses residues 495 to 976, a 482-residue C-terminal deletion with sixteen substituted residues at the junction, which takes the transmembrane helix at residues 538 to 558 and the whole cytoplasmic kinase region and again leaves a truncated extracellular segment. It rises from 0.19 to 0.64 on ipTM and from 0.00 to 0.551 on ipSAE, approaching the threshold without crossing it. CHEK2 removes its FHA regulatory phospho-binding domain together with a polar low-complexity N-terminal segment, residues 1 to 221 (Li et al. 2002), and moves from 0.48 to 0.62 on ipTM while moving from 0.282 to 0.313 on ipSAE. ESR1 removes its N-terminal AF-1 transactivation region, residues 1 to 173, a disordered hormone-coactivator interaction surface (Liu et al. 2024), and moves from 0.48 to 0.60 on ipTM while its ipSAE stays level at 0.802 and 0.804. The ESR1 canonical therefore already holds a confidently predicted interface on the length-corrected metric, and what truncation changes is the whole-chain average. GFER behaves the same way, rising from 0.57 to 0.88 on ipTM while its ipSAE holds at 0.866 and 0.868. Figure 5 places these models beside their canonical counterparts. PRDX5 and FH behave the same way in miniature: both delete a mitochondrial transit peptide and both rise on ipTM, but PRDX5 holds at 0.863 and 0.835 on ipSAE while FH rises from 0.133 to 0.731. Case-level reading of emergence therefore requires the length-corrected metric, and the four cases span the full range from reproduction on both metrics to movement on one alone.

**Figure 5.**
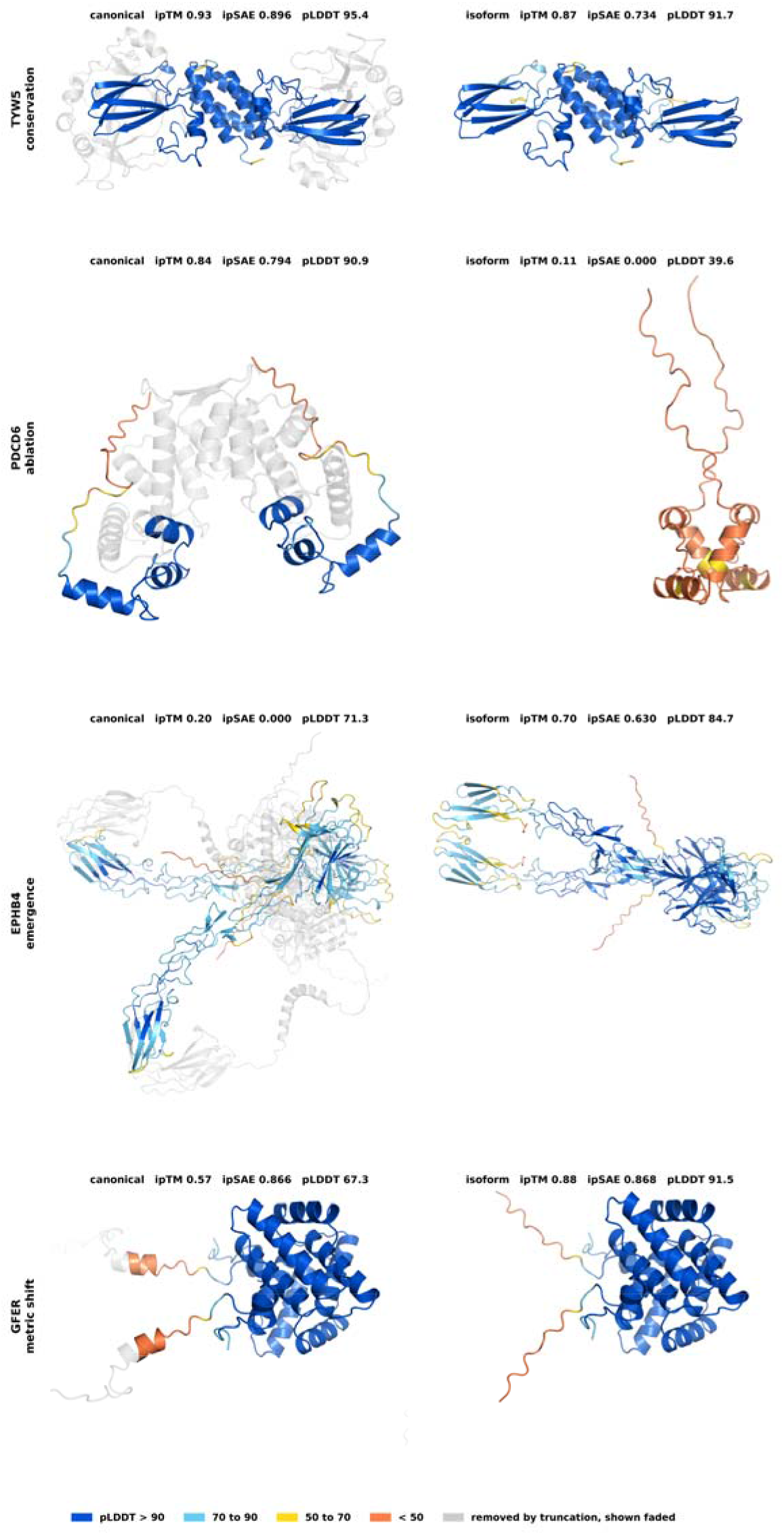
Predicted homodimer models for four representative events, canonical beside truncated isoform. Each row places the predicted canonical homodimer beside the predicted homodimer of its truncated isoform. Within each row the isoform is superposed on its canonical through one chain and the two are drawn at one scale, set by the ordered part of each model, so size and orientation are comparable. Cartoons are coloured by pLDDT, and on each canonical the segment the isoform removes is shown faded. Panel titles give ipTM, ipSAE and mean pLDDT. TYW5 loses 163 N-terminal residues, slightly over half the chain, and both models stay confidently predicted (95.4 and 91.7). PDCD6 loses residues 70 to 191 and the isoform model falls to a mean pLDDT of 39.6, two extended chains in marginal contact rather than an ordered dimer. EPHB4 loses 573 residues, taking the transmembrane helix and the whole cytoplasmic region, and its isoform is predicted more confidently than its canonical, 84.7 against 71.3. GFER loses an 80-residue N-terminal segment whose removal raises mean pLDDT from 67.3 to 91.5 and ipTM from 0.57 to 0.88 while ipSAE holds at 0.866 and 0.868, so the whole-chain average moves while the length-corrected interface score does not.

### Functional diversity and pathway enrichment

We mapped the emergence cohort to the STRING human protein-protein interaction network (Figure S2) and performed pathway enrichment against a custom background of all 4,114 canonical genes. The STRING projection placed 114 of the 188 emergence genes onto an interaction network of 276 edges, of which 72 were high-confidence (combined score ≥ 0.7) and 30 were highest-confidence (≥ 0.9) (Table S8); the mean combined score across the network was 0.60. Degree centrality identified ESR1 (degree 22), MMP2 (22), CASP9 (18), FGF13 (16), NCAM1 (16), CHEK2 (16), ECI2 (14), NT5C1B (14), EPHA2 (14), AK5 (12), PGR (12) and EPHB4 (12) as principal hubs. ESR1, CHEK2, EPHA2 and EPHB4 lay at the connectivity core of the cohort: ESR1 and CHEK2 fell among the top six hubs by degree, EPHA2 in the next tier (degree 14, top nine), and EPHB4 (degree 12) adjacent in the same connected sub-network. The cohort populates a coherent interactome region, even if it does not concentrate in a coherent functional category.

Pathway enrichment for the emergence cohort, computed against a custom background of all 4,114 canonical genes with the statistical domain restricted to annotated genes, returned a single significant term under g:SCS correction at α = 0.05: GO:0006753 nucleoside phosphate metabolic process (p = 0.0379, 24 of 178 annotated query genes; Table S9). Retaining unannotated background genes in the statistical domain admits one further nested term from the same branch, GO:1901293 nucleoside phosphate biosynthetic process (Table S9). The restricted domain is the conservative choice and is the one reported here. The 42 emergence events that reproduce on ipSAE, 36 genes once events sharing a gene are counted once, returned no significant terms, a cohort size at which the absence of terms bounds rather than excludes a functional signal.

The same procedure applied to the complete-ablation cohort was a query of 54 annotated genes beside 178 for the emergence cohort, and returned 14 significant terms (Table S9; Figure 6). These include catalytic activity (42 of 52 genes, p = 9.1 × 10^−7^), identical protein binding (26 of 52, p = 2.7 × 10^−5^), protein homodimerization activity (14 of 52, p = 0.0096) and small molecule metabolic process, together with metabolic pathway terms from KEGG and Reactome. Four of the 14 terms draw their entire intersection from five UDP-glucuronosyltransferase genes at two loci and carry the weight of one observation.

**Figure 6.**
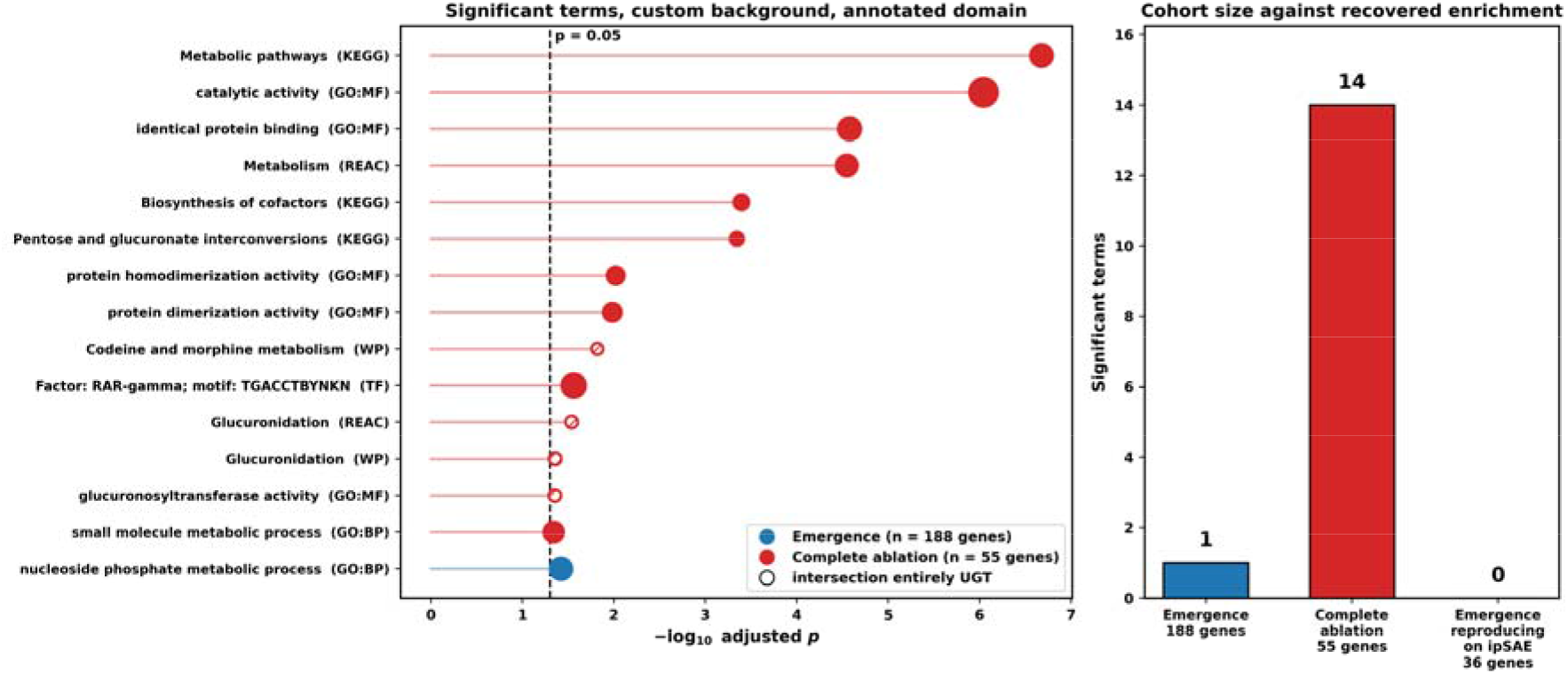
Functional enrichment of the emergence and complete-ablation cohorts. (a) Every term reaching significance under g:SCS correction at α = 0.05 for either cohort, computed against a custom background of all 4,114 canonical genes with the statistical domain restricted to annotated genes. Marker area scales with the number of query genes in the term, and open markers denote terms whose entire intersection is UDP-glucuronosyltransferase genes. The dashed line marks p = 0.05. (b) Number of significant terms against the size of the gene list submitted. The complete-ablation cohort submits 55 genes, of which 54 carry an annotation, and returns 14 terms; the emergence cohort submits 188, of which 178 carry an annotation, and returns 1; the 36 genes behind the emergence events reproducing on ipSAE return none.

The background is itself the canonical set from the homodimer release, so enrichment for protein homodimerization activity within it indicates that proteins whose predicted interface is abolished by truncation are more often annotated homodimers than the remainder of that set. The ablation cohort was drawn from that same background and scored under identical settings, which shows that the procedure recovers functional structure where a cohort carries it. Assessed against that reference, the emergence cohort shows no detectable concentration in any functional category beyond the single nucleotide-metabolism term.

The emergence cohort does not concentrate in any specific functional category, and the structural rules therefore operate across diverse gene-functional contexts and not within any single pathway or family. This functional diffuseness is the natural complement to the structural reproducibility: emergence is governed by topological and biophysical properties of the deletion, namely where the deletion is, and what kind of sequence it removes. The structural grammar is, in this sense, agnostic to gene-functional category, a property that distinguishes it from pathway-specific regulatory mechanisms and places it at the same level of generality as other proteome-scale structural rules such as the correlations between protein size and oligomerization state, between disorder content and protein turnover, between fold class and assembly geometry.

### Conclusions and Potential Implications

We have used the AlphaFold Database homodimer expansion to compare canonical-versus-truncated homodimer architectures across the human proteome. Truncation partitions the landscape into reproducible state-transition classes, and those classes are distinguishable on predicted interface extent as well as on interface confidence: conservation events retain most of the canonical interface, destabilization events roughly two thirds of it, and ablation events none. Interface loss is graded by how much sequence a deletion removes, and position modulates that grading, with C-terminal losses roughly halving the chance of conservation once the deletion exceeds the cohort median. Deletions at either terminus are comparably well tolerated when they are small. Emergence of an interface from a sub-threshold baseline is a narrower phenomenon than a single confidence metric suggests, and the events that reproduce across metrics are identifiable by growth of the predicted interface rather than by the disorder content of the canonical protein.

The conceptual reframing this supports concerns how truncation should be thought about at the structural level. Genetics and transcriptomics treat truncation-bearing splice isoforms as binary perturbations whose principal consequence is loss of function relative to the canonical product. At proteome scale the structural picture is continuous. Truncation acts as a lever across a graded landscape in which deletion magnitude sets the expected loss of interface and deletion position modulates it, so the natural object is a structural rheostat and not a knockout. That reading places truncation beside other proteome-scale structural regularities, among them the relationships between protein size and oligomerization state, between intrinsic disorder content and protein turnover, and between fold class and assembly geometry. It also makes the prior usable: for an unannotated isoform, the fraction of the chain removed and the end it is removed from give a calibrated expectation for whether the homodimer interface survives, before any structure is examined.

The methodological consequence generalizes beyond this dataset. Interface confidence scores based on whole-chain averages respond to chain length and to poorly predicted segments, so a comparison between proteins of different lengths includes a term unrelated to the interface. Where a release distributes a length-corrected metric and counts of interface contacts, a classification should be reported on those as well, and the agreement between them is what licenses a structural reading. The same discipline applies to deletion topology, which exact string matching misassigns whenever splicing leaves novel residues at a junction, and to feature-overlap rates, which need a null matched on window size. Failure modes of this kind are often computable before a score is used, as they are for the gap handling and clade structure behind alignment-based conservation (Karagöl, T. & Karagöl, A. 2026d). Any annotated isoform set can be projected onto its canonical reference layer under this discipline, extending to other oligomeric states, to other species, and to heterodimers, which are outside the present scope and where the biologically important questions about competition between isoform and canonical product lie. Residue-level annotation layers developed for large-scale variant profiling, including co-evolutionary profiles, dynamics-aware evolutionary profiles and directional substitution asymmetry measured from alignments (Karagöl, T. et al. 2025; Karagöl, T. & Karagöl, A. 2026b; Karagöl, A. & Karagöl, T. 2026b), map naturally onto the deletion windows and would identify which removed residues account for a lost interface.

ESR1, CHEK2, EPHA2 and EPHB4 lie at the connectivity core of the emergence cohort and each illustrates a distinct mode, and the subset that reproduces across independent interface metrics is the natural starting set for biochemical validation and for all-atom simulation of the kind applied to a monoamine transporter truncation isoform (Karagöl, T. & Karagöl, A. 2026c). All of these are predicted-structure observations, and whether any specific predicted homodimer is realized in cells is a separate question.

The analytical unit is a predicted homodimer and the quantities reported are interface confidence scores and predicted interface extent, so binding affinity, stoichiometry, half-life and cellular complex formation are not addressed. AlphaFold identifies homodimers less reliably than heterodimers, at a true-positive rate of 18% for homodimers and 63% for heterodimers at a 1% false-positive rate (Danielsson and Elofsson 2025), so the quantities reported here are differences between two predictions of the same protein from one method and not absolute calls on either. Only homodimer-versus-homodimer comparisons are modeled, so a gene whose homodimer is sub-threshold may still form a competent heterodimer, and the rules hold as statistical statements about populations and not as deterministic predictions for individual molecules.

Truncation, read at the level of predicted homodimer architecture across the human proteome, is a graded and topology-asymmetric structural lever whose statistics follow the physics of interfaces. Deletion size and deletion position predict the fate of a homodimer interface across the human proteome, for any isoform, before experiment.

## Methods

### Datasets and canonical homodimer reference layer

The canonical human homodimer reference layer was obtained from the recent AlphaFold Protein Structure Database (AFDB) expansion (processed from an initial release manifest of approximately 31 million predicted structural complexes), in which predicted homodimer structures and AlphaFold-Multimer interface confidence scores (interface predicted Template Modeling, ipTM) were generated across the human proteome in collaboration with NVIDIA (Varadi et al. 2024; Abramson et al. 2024; Han et al. 2026). Annotated truncated isoforms for each canonical UniProt entry were obtained from UniProt (UniProt Consortium 2025), retaining alternative-splicing, alternative-promoter, and post-translational-truncation isoform classes and excluding fragment-only entries. Each canonical UniProt accession was paired with every annotated truncated isoform of the same gene. All canonical and isoform sequences were retrieved in FASTA format. No structure prediction was performed for this study. Both the canonical and the truncated-isoform homodimer ipTM values were read from the same AFDB release manifest, which already contains AlphaFold-Multimer homodimer predictions for isoform-suffixed accessions alongside canonical ones (Evans et al. 2022; Han et al. 2026). Comparability between the two therefore follows from their having been generated by one analysis within one release. We retained interface predicted Template Modeling (ipTM) as the primary confidence metric throughout the analysis.

### Interface metrics

The release manifest distributes, for every predicted complex, the interface predicted Template Modeling score (ipTM), the interface Scored Alignment Error in both chain directions (ipSAE), pDockQ and pDockQ2, the Local Interaction Score (LIS), the number of inter-chain interactions, and the counts of residues inside the interface distance cutoff in each direction. We report ipTM throughout and took ipSAE_min, the lower of the two chain directions, as the primary metric, matching the release’s own definition of its high-confidence set and of its confidence tiers. For homodimers the two chain directions of ipSAE agree closely (Pearson r = 0.9994). ipSAE_min is zero for 57.2% of canonicals, meaning that no residue pair passes the alignment-error cutoff, so all comparisons on it use medians and rank statistics. We took interface extent as the sum of the two directional counts of residues inside the interface distance cutoff. The count of inter-chain interactions scales with chain length (Spearman rho = 0.173 against the length change) and was not used as an interface measure.

### Gene-centric canonical-isoform pairing within base-accession families

We retained Homo sapiens entries with a non-null AlphaFold-Multimer ipTM score and recovered the corresponding amino-acid sequence for each accession from the AFDB-distributed FASTA archive (UniProt Consortium 2025). The pairing follows the gene-centric isoform-map approach we previously applied to glutamate transporters (Karagöl, A. et al. 2024), extended here to the full proteome. To assemble (canonical, isoform) pairs strictly from same-locus protein products, accessions were grouped by base accession, defined as the UniProt primary accession after stripping any trailing isoform, so that all members of a group share locus assignment by construction and cannot be contaminated by entries from other loci that merely share a gene symbol. Within each family the reference member was the accession equal to the base accession or ending in -1, taking the higher ipTM where both were present. UniProt’s displayed canonical sequence is not always the -1 isoform. In 19 of the 5,168 pairs this rule selected a longer non-displayed isoform as the reference and placed the displayed canonical in the truncated position; those pairs remain valid longer-versus-shorter comparisons and are identified in Table S1.

The non-canonical members were retained as candidate truncated isoforms only if they passed a subsequence test against the reference. Candidates occurring verbatim within the reference were accepted. Otherwise the longest matching block between the two sequences was computed with the Python difflib SequenceMatcher, and the candidate was accepted when that block covered at least 95% of its length. We discarded members failing the test as non-truncations. An isoform that skips an internal exon and leaves two shorter matching blocks fails this test, so internal exon-skipping events are under-represented in the cohort and the deletions analysed here are predominantly terminal. SequenceMatcher was called with its default settings, which enable the autojunk heuristic; for a candidate of 200 residues or more this restricts which residue types may seed a match, although the seed is then extended through all matching positions. With the heuristic disabled, 400 of the accepted non-subsequence pairs return identical verdicts. This is a near-subsequence criterion and not an exact one. Of the 5,159 pairs whose two sequences could be re-retrieved from UniProt at the lengths recorded in Table S1, 2,743 are exact subsequences of their reference and 2,416 are not. The latter match their reference across at least 95% of their length with a median of 7 residues outside that block, with a 90th percentile of 20. Those pairs are truncations accompanied by local sequence change, and are not pure deletions.

We applied no reviewed-versus-unreviewed filter, because the AFDB homodimer layer is populated against UniProtKB in its full scope and a Swiss-Prot-only restriction would have discarded predictions the manifest already provides. Base-accession grouping operates within a single UniProtKB entry, pairing a canonical sequence only with isoform-suffixed accessions of that same entry, and never merges two entries. Each unreviewed accession therefore forms its own family instead of being attached to a reviewed one.

### Tier stratification and four-class state-transition assignment

We partitioned canonical homodimer predictions into three confidence tiers based on canonical ipTM: High (canonical ipTM ≥ 0.8; n = 234), Moderate (0.6 ≤ canonical ipTM < 0.8; n = 354), and Low (canonical ipTM < 0.6; n = 4,580). The 0.8 boundary is a widely used AlphaFold-Multimer high-confidence interface cutoff. The 0.6 boundary is the lower bound applied by the mapping analysis and is reported here as implemented; it is not drawn from a published convention. Each (canonical, isoform) pair was then assigned to one of four state-transition classes based on the joint canonical and isoform tier classification: complete interface ablation, structural conservation, partial destabilization, and truncation-associated emergence. The strict gain class comprises all Low→Moderate, Low→High, and Moderate→High transitions (n = 208 in the present dataset), and is also the cohort used for downstream biophysical and annotation-level analysis.

### Deletion window derivation and topology classification

Deletion windows in canonical numbering were derived for every pair by global pairwise alignment of canonical against isoform sequence with Biopython’s PairwiseAligner (Cock et al. 2009) under BLOSUM62, gap-open -11, gap-extend -1, and end-gap penalties set to zero so that terminal deletions are not penalized. We extracted canonical positions aligned to gaps in the isoform as deletion intervals, and non-contiguous intervals within one isoform were retained separately. We assigned truncation topology from the span of those intervals, with a terminal tolerance applied at each end to absorb novel residues introduced at splice junctions. The tolerance was swept from 0 to 30 residues; the proteome-wide C-terminal share moves from 42.7% to 43.3% across that range. Exact prefix and suffix matching was not used, because a C-terminal truncation that leaves novel junction residues before the stop codon produces a window that stops short of the final residue, which a strict terminal test reads as an internal deletion.

### UniProt feature mining

For each unique canonical UniProt accession in the emergence cohort, the full UniProt JSON entry was retrieved from UniProt and parsed for curated structural features. We retained fourteen feature classes: Domain, Region, Motif, Repeat, Compositional bias, Coiled coil, Zinc finger, DNA binding, Active site, Topological domain, Transmembrane, Signal peptide, Propeptide, and Binding site. Each feature was intersected with each deletion interval in canonical numbering; overlap of ≥ 1 residue was scored as a hit, with the overlap tagged as full or partial based on whether the entire feature was contained within the deletion. Lexical tagging applied two regular-expression matchers to feature type and description fields: an inhibitory pattern (autoinhibit*, inhibitory, pseudo-substrate, regulatory, pre-segment, prodomain, propeptide, intramolecular-lock, masking, occluding) and a disorder / compositional-bias pattern (disordered, low-complexity, polar, acidic, basic, Gly-rich, Pro-rich, Ser-rich, polyampholyte, compositional bias).

### Random-window null for feature deletion

Rates of overlap between deletion windows and curated UniProt features were compared against a null in which a window of the same length is placed uniformly at random within the same canonical sequence, with 2,000 draws per event and a fixed seed. The null was computed separately for any curated feature, for the disorder and compositional-bias classes, and for features whose type or description matches inhibitory, regulatory or pseudo-substrate language.

### Disorder and binding-propensity analysis

We took canonical disorder as the fraction of residues with pLDDT below 50, read from the AFDB prediction metadata for every canonical accession and not recomputed from the model files. Agreement with per-model computation is exact on the profiled cohort (Pearson r = 1.0000). Because disorder depresses interface confidence and the emergence and ablation cohorts are selected at opposite ends of the canonical confidence range, the comparison between them is confounded with the selection rule. The comparison reported as primary therefore holds the canonical stratum fixed, setting the emergence cohort against all other pairs whose canonical ipTM falls below the same threshold, and is repeated within narrow canonical ipTM bins. Effect sizes are Cohen’s d from pooled standard deviation and significance is from the two-sided Mann-Whitney U test. Peak per-residue binding propensity was computed with AIUPred (Erdős and Dosztányi 2024) as the highest score anywhere in a sequence, and its dependence on sequence length was evaluated across the profiled cohorts by Spearman rank correlation.

### Protein-protein interaction network projection

Gene symbols in the emergence cohort were mapped to STRING identifiers using the STRING website (string-db.org v12.0, organism Homo sapiens, taxonomy 9606) (Szklarczyk et al. 2023). The resulting network was retrieved as a TSV containing protein-protein interaction edges with combined score and individual evidence channels (neighborhood, gene fusion, phylogenetic co-occurrence, homology, co-expression, experimentally determined, database-annotated, automated text-mining). We retained edges at the default STRING confidence threshold (combined score ≥ 0.4); high-confidence (≥ 0.7) and highest-confidence (≥ 0.9) subsets. Node degree centrality was computed as the count of edges incident on each node.

### Collapsing shared splicing events

Events sharing one splicing event across paralogues were identified by grouping events of identical deletion length and aligning their deleted segments, since paralogues that share only part of their exon structure are not grouped by whole-chain identity. We collapsed pairs whose deleted segments align above an identity threshold to one event. The threshold was swept from 0.80 to 0.99 without changing the groupings recovered.

### Pathway enrichment analysis

Functional enrichment was performed with g:Profiler (Kolberg et al. 2023), database version e114_eg62_p19_9dc92ac8, using a custom background of canonical gene symbols comprising every gene that entered the AFDB-Multimer scoring analysis. We restricted the statistical domain to annotated genes within that background, so that genes with no annotation did not enter the domain size. We queried three cohorts under identical settings: the emergence cohort, the complete-ablation cohort, and the subset of emergence events reproducing on ipSAE. The full default set of g:Profiler data sources was queried jointly, of which Gene Ontology, KEGG, Reactome, WikiPathways and TRANSFAC returned terms. Multiple-testing correction used g:SCS at α = 0.05, which accounts for the dependence between nested ontology terms. We applied no term-size cap. Enrichment computed against the same background with unannotated genes retained in the statistical domain is reported alongside the primary result in Table S9.

### Comparison with deposited structures

Canonical accessions were mapped to Protein Data Bank entries through their UniProt cross-references, and deposited chains to UniProt accessions through SIFTS (Dana et al. 2019). Deposited biological assemblies were read from PDBe (Armstrong et al. 2020). For each assembly we summed the number of copies of the polypeptide entities whose chains map to the canonical accession, and recorded the largest such count across all assemblies of all entries for that accession. Two or more copies was taken as deposited evidence for a homomer, and assemblies containing no other polypeptide were recorded separately as the stricter criterion. Rates were compared between confidence tiers with the two-sided Fisher exact test.

### Statistical analyses

Cohort comparisons used the two-sided Mann-Whitney U test with Cohen’s d from pooled standard deviation. Associations between continuous quantities used Spearman rank correlation, chosen because ipSAE has a large point mass at zero. Comparisons across the three High-tier outcome classes used the Kruskal-Wallis test. Confidence intervals on proportions are bootstrap percentile intervals from 4,000 resamples with a fixed seed. Thresholds transferred between metrics on different scales were matched by quantile against the ipTM tiers and not transferred directly. All statistics were computed with scipy.stats, numpy and pandas (Virtanen et al. 2020; Harris et al. 2020; McKinney 2010).

## Supplementary information

Supplementary_tables.xlsx, holding Supplementary Tables S1 to S13, and Supplementary_information.pdf, holding Supplementary Figures S1 and S2.

## Data availability statement

The data and custom code supporting this article are freely available to the scientific community, in our public GitHub repository at https://github.com/karagol-taner/truncation-structural-grammar. Publicly available datasets and web tools analyzed in this study can be accessed at their respective primary repositories. Canonical homodimer predictions and AlphaFold monomer models were retrieved from the AlphaFold Protein Structure Database (https://alphafold.ebi.ac.uk/). Protein sequence data and curated structural feature annotations were obtained from UniProt (https://www.uniprot.org/). Protein-protein interaction network projections were generated with STRING (https://string-db.org/). Pathway enrichment analyses were conducted with g:Profiler (https://biit.cs.ut.ee/gprofiler/). An earlier version of this work is available as a preprint (https://doi.org/10.64898/2026.05.06.723091). Any additional data not included in the main text or supplementary materials are available from the corresponding authors upon reasonable request.

## Ethics approval

Ethics approval was not required for this computational study as it did not involve animal subjects, human participants, and identifiable data.

## Consent to participate

Not applicable.

## Consent for publication

Not applicable.

## Author contributions

A.K. and T.K. contributed equally to this work.

## Declaration of Generative AI and AI-Assisted Technologies in the Manuscript Preparation Process

During the preparation of this work, the authors used Claude Opus 5 (Anthropic) for drafting and for streamlining analysis code. The authors reviewed and edited the output as needed and take full responsibility for the content of the published article.

