## Supplementary_information for "Structural grammar of truncation across the human homodimer landscape"

### Table of Contents


| | | |
|---|---|---|
| **1.** | Figure S1. Controls for the feature-deletion rate and for the length-dilution model. | Pg. 3 |
| **2.** | Figure S2. STRING protein-protein interaction network of the emergence cohort. | Pg. 4 |
| **3.** | Table S1. Main landscape dataset: every pair on four interface metrics, with realigned deletion windows | xlsx |
| **4.** | Table S2. Aggregated topology against outcome class | xlsx |
| **5.** | Table S3. Complete-ablation cohort | xlsx |
| **6.** | Table S4. Partial-destabilization cohort | xlsx |
| **7.** | Table S5. Emergence cohort, with reproduction on ipSAE flagged | xlsx |
| **8.** | Table S6. Canonical pLDDT-defined disorder, all canonicals with cohort labels | xlsx |
| **9.** | Table S7. Deletion windows and cleavage-site overlap in the emergence cohort | xlsx |
| **10.** | Table S8. STRING protein-protein interactions | xlsx |
| **11.** | Table S9. Gene Ontology and pathway enrichment for the emergence and complete-ablation cohorts against a custom background of all canonical genes. The 36-gene subset of emergence events reproducing on ipSAE was tested under identical settings and returned no significant terms, so it contributes no rows | xlsx |
| **12.** | Table S10. Agreement between the four interface metrics and class transitions | xlsx |
| **13.** | Table S11. Matched controls: random-window null, disorder strata, interface retention, bootstrap intervals | xlsx |
| **14.** | Table S12. Shared splicing events across paralogues, and topology under each assignment | xlsx |
| **15.** | Table S13. Deposited-structure state of the canonical proteins: largest number of copies in any deposited biological assembly, for every canonical carrying a Protein Data Bank entry | xlsx |


Supplementary Tables S1 to S13 are supplied as sheets of the workbook Supplementary_tables.xlsx.

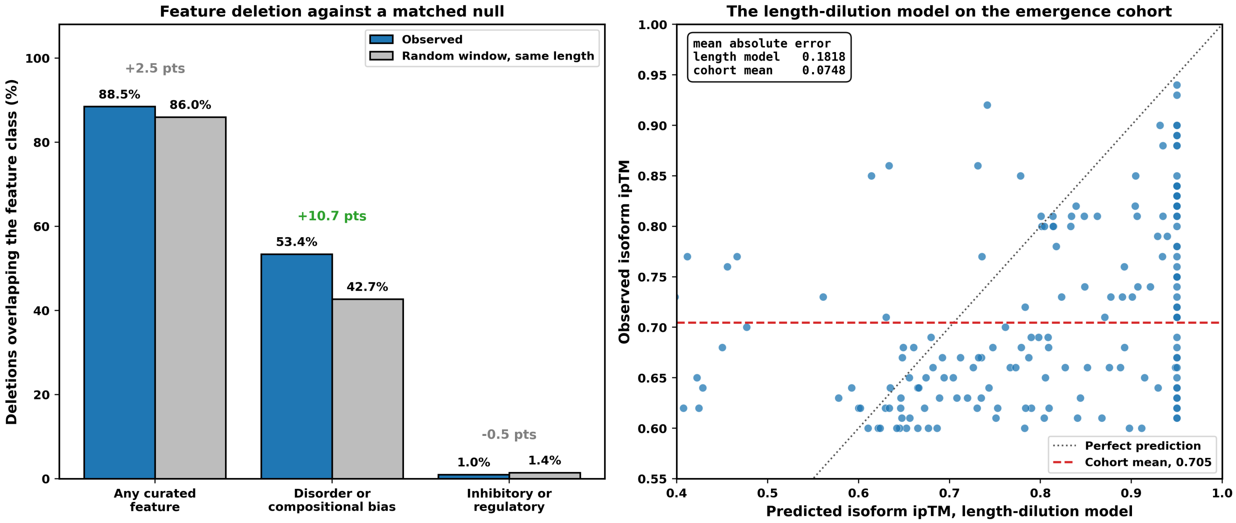


**Figure S1. Controls for the feature-deletion rate and for the length-dilution model.**
(a) Fraction of emergence-cohort deletions overlapping each feature class, against a null in which a window of the same length is placed uniformly at random within the same canonical sequence, 2,000 draws per event. The difference in percentage points is printed above each pair. (b) Observed isoform ipTM against the prediction of a model that scales canonical ipTM by the ratio of canonical to isoform length and caps the result. The dashed line marks the cohort mean. The model reproduces the cohort mean while carrying a per-pair error 2.4 times that of predicting the cohort mean for every pair.

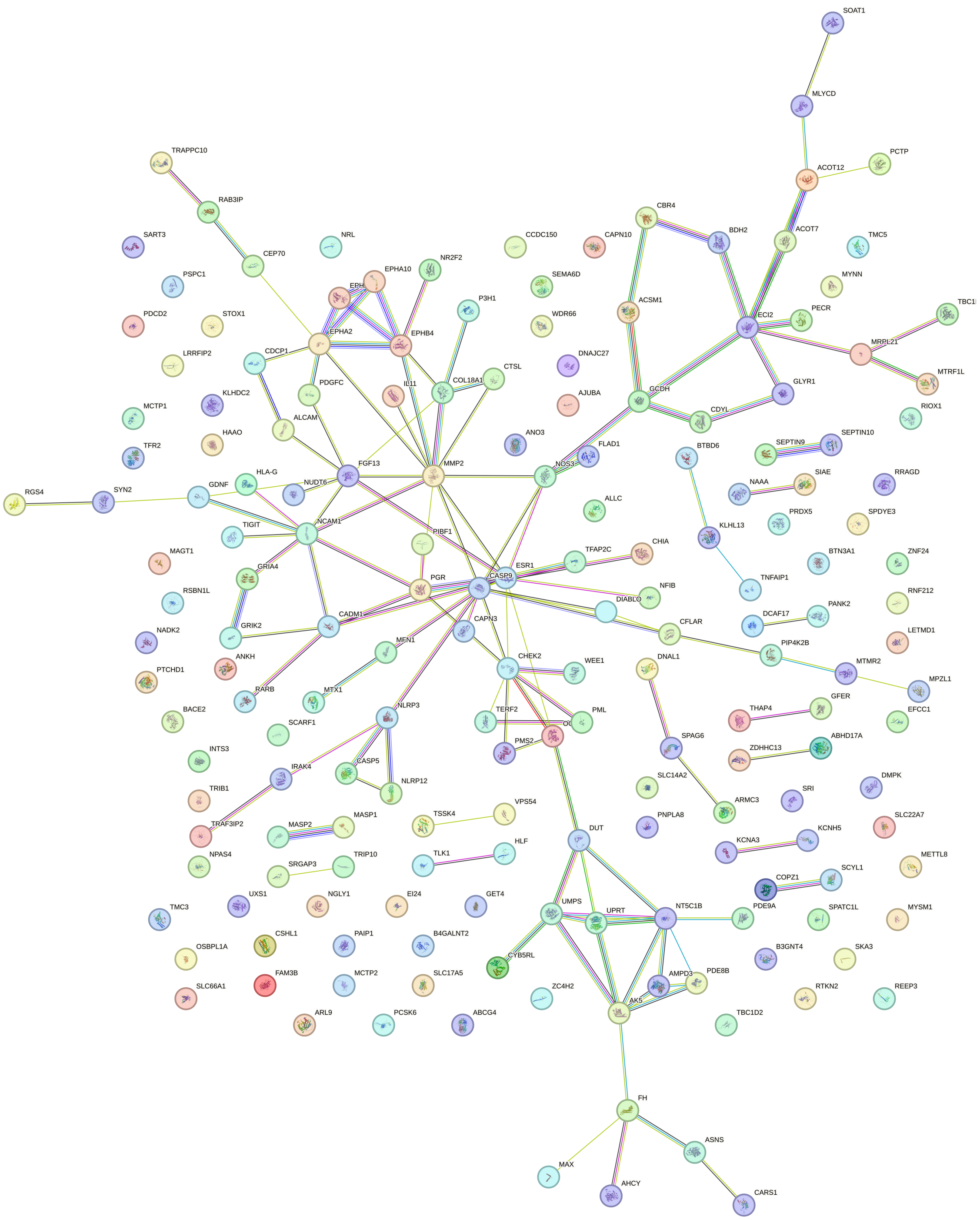

**Figure S2. STRING protein-protein interaction network of the emergence cohort.**
STRING v12.0 (Homo sapiens, taxId 9606) maps 114 of 188 emergence genes onto a network of 276 edges, of which 72 are high-confidence (combined score 0.7 and above) and 30 highest-confidence (0.9 and above); the mean combined score is 0.60. Node size, degree centrality. Principal hubs: ESR1 (22), MMP2 (22), CASP9 (18), FGF13 (16), NCAM1 (16), CHEK2 (16), ECI2 (14), NT5C1B (14), EPHA2 (14), AK5 (12), PGR (12), EPHB4 (12). Edge width, STRING combined score.
